# High syphilis seropositivity in European brown hares, Lower Saxony, Germany

**DOI:** 10.1101/2020.01.05.893719

**Authors:** Linda Hisgen, Lena Abel, Luisa K. Hallmaier-Wacker, Simone Lueert, Ursula Siebert, Marcus Faehndrich, Egbert Strauss, Ulrich Voigt, Markéta Nováková, David Šmajs, Sascha Knauf

**Affiliations:** Georg-August-University, Goettingen, Germany; German Primate Center, Goettingen, Germany; University of Veterinary Medicine Hanover - Foundation, Hanover, Germany; Masaryk University, Brno, Czech Republic

**Keywords:** *Treponema paraluisleporidarum*, Lagomorpha, *Lepus europaeus*, Europe, syphilis, serology

## Abstract

*Treponema paraluisleporidarum* infects lagomorphs and is a close relative of the human syphilis-bacterium *Treponema pallidum*. There is paucity of information on the epidemiology of hare syphilis and its relationship to rabbit- and human-infecting *Treponema*. We have found a high seropositivity (405/734) for *Treponema paraluisleporidarum*-infection in hares of Lower Saxony, Germany.

## Text

In humans, *Treponema pallidum* is the causative agent of syphilis (subsp. *pallidum* (*TPA*)), yaws (ssp. *pertenue*) and bejel (ssp. *endemicum*). Although its closest genetic relative, *Treponema paraluisleporidarum* ecovar Cuniculus (*TP*eC), which infects rabbits, is believed to be apathogenic to humans [1], its genome differs in less than 2% from *TPA* [2]. Experiments have shown that the hare-infecting ecovar Lepus (*TP*eL) leads to clinical disease in both, rabbit and hare, whereas *TP*eC causes only seroconversion in the European brown hare (*Lepus europaeus*, EBH) [3].

In this study, we aimed to investigate the presence of *TP*eL in wild EBHs and hypothesized, that the disease is widespread among the hare population of Lower Saxony (LS), Germany. We used serological tests to demonstrate the presence of antibodies against *TP*eL. EBHs are rarely reported with clinical lesions such as skin ulcerations and crusting of the anogenital and facial region [4]. However, similar to what is seen in human syphilis, infection elicits a pronounced serological response in its lagomorph host [5]. Since bacterial clearance is not reported, and animals in the wild are not treated with antibiotics, EBHs with antibodies against *TPeL* were considered to be infected. We used the Epitools FreeCalc software (https://epitools.ausvet.com.au/freecalctwo) to calculate the required sample size to demonstrate population freedom from disease, using imperfect tests and allowing for small populations [6]. Population size was used as shown in Table S1. Test sensitivity (0.97) and specificity (0.92) were set according to the test performance reported in Knauf et al. 2012 [7]. Recent studies report a prevalence in EBHs from 33.0 to >74%, [8, 9]. We, therefore, calculated sample size with a design prevalence of 30%. The desired type I and type II error were each set to 0.05. All other parameters were kept at default settings.

Based on our calculated sample size (711), we used a total of 734 serum samples collected from shot hares between 2007 and 2018 from several hunting areas in LS (Technical Appendix Table S1 and S2). Corresponding age data were used to group the sampled EBHs into age classes.

All serum samples were screened utilizing the Serodia TP-Particle Agglutination assay (Serodia® TP-PA, Fujirebio Diagnostics Inc., Malvern, PA, USA). This assay does not require species-specific secondary antibodies, which makes it particularly suitable for the use in wildlife. In the presence of reactive antibodies, the antigen-coated gelatine particles are cross-linked, which results in an even-coated bottom of the reaction well. None of the samples generated positive reactions with galatine particles without coated antigen on their surface (control). A statistically representative number of randomly selected samples (233/734) were additionally confirmed using a fluorescent treponemal antibody absorption test (FTA-ABS, Mast Diagnostica, Reinfeld, Germany) that was modified using a fluorescein isothiocyanate-conjugated secondary anti-rabbit immunoglobin G (Sigma-Aldrich, Cat. no: F9887).

The Serodia TP-PA test became reactive in 55% (405/734) of the tested animals. 215 out of 374 female (57%) and 167 out of 325 (51%) male EBHs tested positive. In subadult EBHs, only 36/267 (13%) showed a positive reaction, whereas 339/417 (81%) of the adult EBHs had cross-reactive antibodies. In 34 animals, the gender was unknown and in 49 animals no age class could be determined. All animals tested with the FTA-ABS assay (233) showed the same result as in the TP-PA test. Figure 1 illustrates the regional rate of positive tested EBHs. Differences in sample size prevented us from reporting the prevalence, however, serological data show that infection is widespread. Using a two-tailed Fisher-exact test and considering p≤0.05 statistically significant, our data show no significant difference of infection between sex (odds ratio: 0.79 [95% Cl 0.58-1.07], p=0.1283), but a significant difference based on age (odds ratio: 0.03 [95% CI 0.02-0.05], p<0.0001) with a lower seropositivity among young hares. The latter argues for a sexual transmission mode, as reported for human and rabbit syphilis [10]. Clinical manifestations have been described to occur in about 1% of the infected wild EBHs [4] and were not seen in any of the examined hares in this study. This is likely attributed to the low number of animals that were prospectively sampled (19) and which were clinically examined.

Nevertheless, the high number of serologically positive, hence naturally infected, EBHs in LS warrants further investigations to perform molecular characterization of hare-infecting strains and subsequent genome sequencing. Currently, *TP*eC strain Cuniculi A (GenBank Accession no.: NC_015714.1) is the only complete genome of lagomorph-infecting *T. paraluisleporidarum*. Since the loss of human-pathogenicity is thought to be associated with deletions in the *tprF*, *G* and *I* gene [2], it will be important to investigate whether hare-infecting strains have the same genome deletions, to rule out zoonotic potential of these strains. The close genetic relationship of lagomorph-infecting treponemes to human-infecting *T. pallidum* provides a unique opportunity to understand the evolution of syphilis and its virulence factors that code for human-pathogenicity.

**Figure 1.**
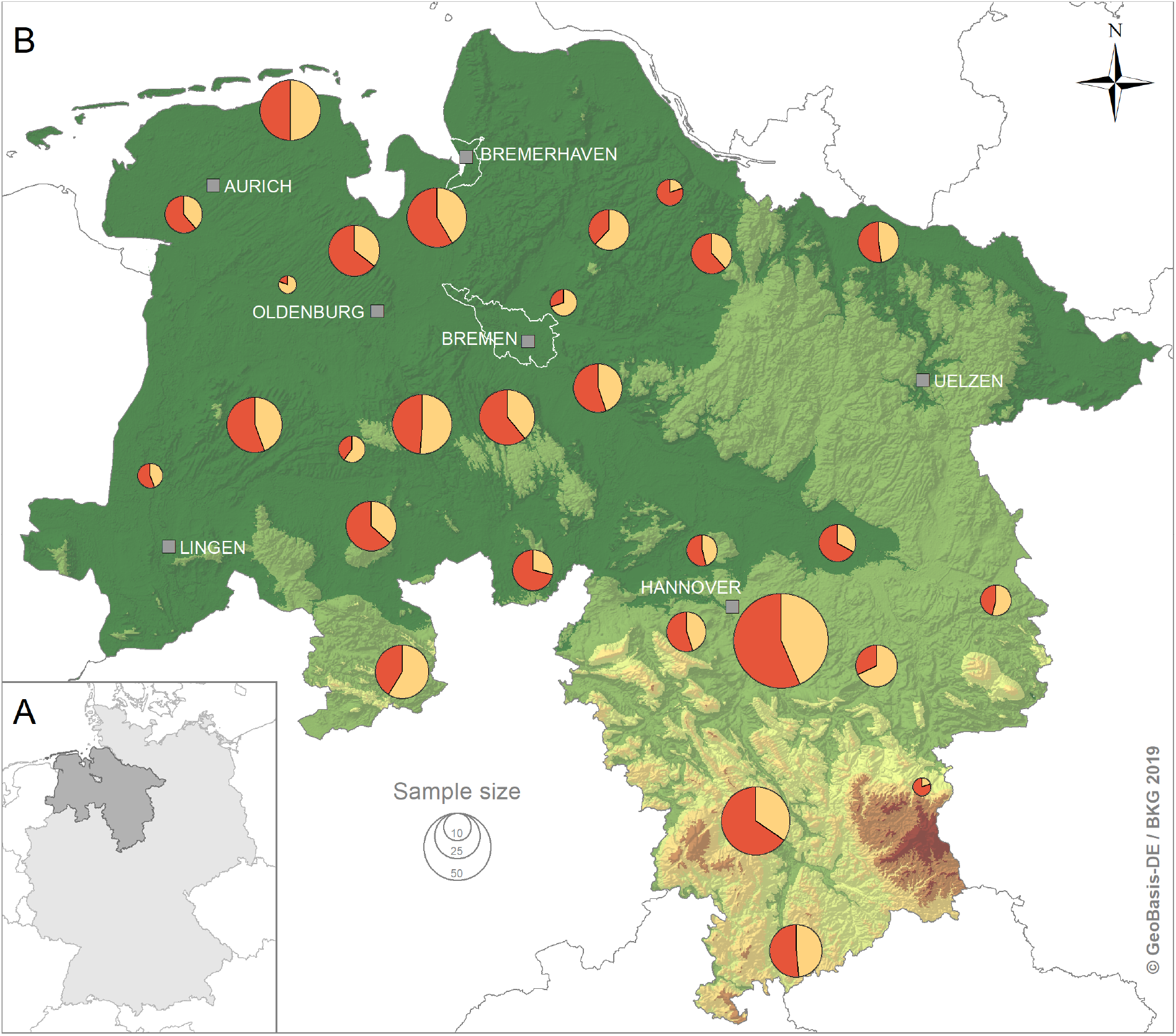
**(A)** The location of Lower Saxony within Germany. **(B)** A map of Lower Saxony and its neighbouring federal states, showing where EBHs were sampled. Circle size represents the sample size; red indicates the proportion of serological positive animals; yellow indicates the proportion of serological negative animals. The map was created using ArcMap version 10.6.1 (ESRI, Redlands, CA, USA; map source: © GeoBasis-DE/BKG 2019).

## Supporting information

Technical Appendix

## Acknowledgements

We would like to acknowledge the Hunting Association of Lower Saxony (Landesjaegerschaft Niedersachsen) for the consent to the project as well as all hunters for providing samples and supporting the study.

This study was funded by the German Research Foundation (DFG KN1097/7-1 to SK) and partly from the Czech Science Foundation (GC18-23521J to DŠ). Sampling was furthermore financially supported by the Ministry of Food, Agriculture and Consumer Protection of Lower Saxony (http://www.ml.niedersachsen.de; to UV). The funders had no role in study design, data collection and analysis, decision to publish or the preparation of the manuscript.

