## Supplementary material for "High syphilis seropositivity in European brown hares, Lower Saxony, Germany": Technical Appendix

### Ethical Statement

No live hares were handled and no hares were purposely shot for this study. Hare samples originated exclusively from legally hunted hares in Lower Saxony, Germany. Ethical clearance for sample collection was thus not required.

**Table S1. Population size data and resulting sample sizes**. Population size is based on wildlife survey data of Lower Saxony [11]. For simplification, calculations assumed that hare populations were stable throughout the study period. All other parameters to calculate the sample size were set as described in the article file. Although the required sample size was not reached in 19/32 geo-groups, only one (geo-group 30) remained without positive tested samples. NA=not applicable. *Some of the hunting districts were samples in multiple years. In these cases, the total number of reports was greater than the number of hunting districts. **Population size data were not available for the year of sampling (2018). The estimates are therefore averaged from data reported in 2007 and 2009.

| Geo-Group | N population size reports (n hunting districts)* | Estimated overall hare pop (n hares) | Median of population size (Range) | Required sample size at 30% design prevalence | Tested samples which are included into the study (n TPeL positive) |
|---|---|---|---|---|---|
| 1 | 32 (8) | 984 | 105.5 (169) | 24 | 41 (23) |
| 2 | 1 (1) | 40 | NA | 21 | 5 (1) |
| 3 | 1 (1) | 173 | NA | 23 | 34 (14) |
| 4 | 3 (2) | 214 | 423.5 (579) | 24 | 36 (22) |
| 5 | 1 (1) | 225 | NA | 23 | 31 (20) |
| 6 | 4 (4) | 236 | 58.0 (30) | 24 | 37 (25) |
| 7 | 4 (5) | 266 | 65.5 (45) | 24 | 52 (34) |
| 8 | 2 (2) | 124 | 62.0 (36) | 23 | 30 (15) |
| 9 | 1 (1) | 101 | NA | 23 | 41 (20) |
| 10 | 1 (1) | 108 | NA | 23 | 10 (3) |
| 11 | 1 (1) | 360 | NA | 24 | 9 (5) |
| 12 | 4 (1) | 275 | 275.0 (100) | 24 | 42 (21) |
| 13 | 1 (1) | 150 | NA | 23 | 22 (7) |
| 14 | 2 (2) | 196 | 98.0 (44) | 23 | 21 (8) |
| 15 | 1 (1) | 95 | NA | 23 | 5 (4) |
| 16 | 2 (1) | 207 | 207.0 (286) | 24 | 30 (19) |
| 17 | 1 (1) | 67 | NA | 23 | 13 (6) |
| 18 | 2 (1) | 986 | 985.0 (29) | 24 | 21 (13) |
| 19 | 1 (1) | 137 | NA | 23 | 7 (4) |
| 20 | 2 (2) | 171 | 85.5 (45) | 23 | 3 (2) |
| 21 | 2 (1)** | 128 | 127.5 (5) | 23 | 29 (16) |
| 22 | 1 (1) | 36 | NA | 21 | 18 (12) |
| 23 | 2 (1) | 113 | 113.0 (26) | 23 | 20 (11) |
| 24 | 1 (1) | 180 | NA | 23 | 21 (15) |
| 25 | 2 (2) | 194 | 97.0 (98) | 23 | 13 (7) |
| 26 | 1 (1) | 200 | NA | 24 | 10 (8) |

| | | | | | |
|---|---|---|---|---|---|
| 27 | 1 (1) | 485 | NA | 24 | 18 (11) |
| 28 | 2 (1) | 340 | 340.0 (160) | 24 | 41 (24) |
| 29 | 2 (1) | 52 | 52.0 (6) | 22 | 14 (4) |
| 30 | 1 (1) | 85 | NA | 22 | 3 (0) |
| 31 | 1 (1) | 110 | NA | 23 | 36 (20) |
| 32 | 1 (1) | 85 | NA | 22 | 21 (11) |
| **Total** | **82** (50) | **6,995** | **NA** | **717** | **734** (405) |

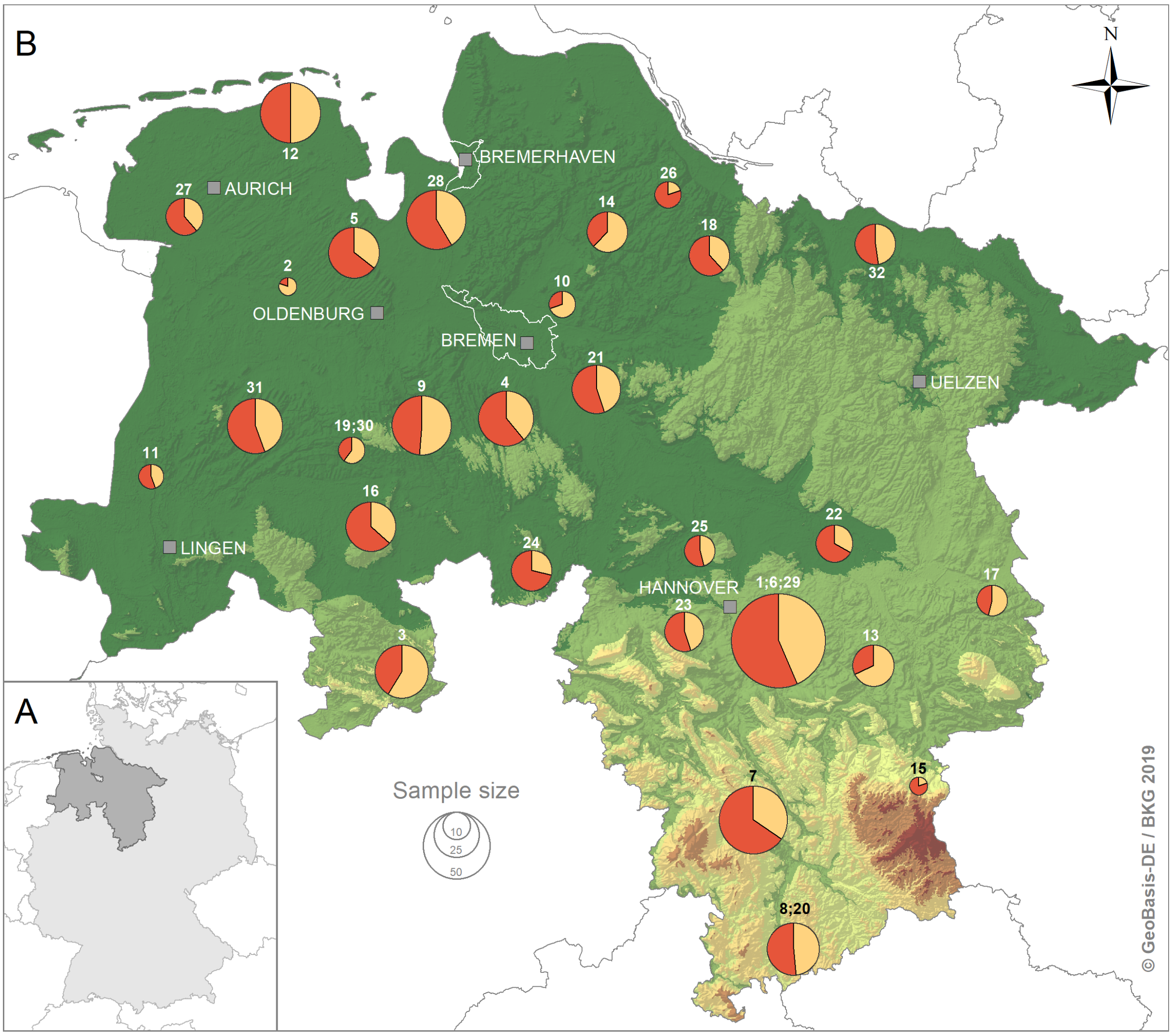


**Figure S1. The figure corresponds to Figure 1 and includes the geo-groups listed in Table S1. (A)** The location of Lower Saxony within Germany. **(B)** A map of Lower Saxony and its neighbouring federal states, showing where EBHs were sampled. Circle size represents the sample size; red indicates the proportion of serological positive animals; yellow indicates the proportion of serological negative animals. Geo-group 19 and 30, 1, 6 and 29 as well as 8 and 20 have been combined for better visualization of the sampling areas. The map was created using ArcMap version 10.6.1 (ESRI, Redlands, CA, USA; map source: © GeoBasis-DE/BKG 2019).

**Table S2. Summary of the serological result according to age and sex**. Serological positive were all animals that reacted in both TP-PA and FTA-ABS test; TP negative are animals, that are negative in both tests.

| | **Sub-adult** | | **Adult** | | **Unknown sex** | | **Unknown age** | | **Unknown sex and age** | |
|---|---|---|---|---|---|---|---|---|---|---|
| | **Male** | **Female** | **Male** | **Female** | **Sub-adult** | **Adult** | **Male** | **Female** | | **Sub-total** |
| **Sero-positive** | 17 | 14 | 141 | 181 | 5 | 17 | 9 | 20 | 1 | 405 |
| **Sero-negative** | 116 | 105 | 32 | 45 | 10 | 1 | 10 | 9 | 1 | 329 |
| **Total** | 133 | 119 | 173 | 226 | 15 | 18 | 19 | 29 | 2 | 734 |

### Age estimation

EBHs were grouped according to different age-related parameters. The primary parameter that was used to group animals into the class ‘subadult’ or ‘adult’ was the weight of the dried eye-lens [12]. In cases where the lens weight was not available, e.g. when both eyes were damaged through munition, animals were classified according to the ossification status of the epiphyseal cartilage of the ulna [13].

*Dried eye-lens weight*

Eye-lenses of shot hares corresponding to the serum samples were collected and processed as described in Suchentrunk et al. 1991 [12]. Briefly, desiccation of lenses was achieved by heating them at 100°C for 24 hours in an ordinary dry cabinet. An analytic scale was used to weight the dry lenses to the nearest 1 mg. Animals with a dried eye-lens weight of greater 275.0 mg were considered adults.

*Ossification of the epiphyseal cartilage*

The ossification status of the epiphyseal cartilage of the ulna can be palpated. The different stages of ossification were used to group EBHs into 'subadults' and 'adults'. Since complete epiphyseal ossification of the ulna is reached after seven to nine months [13], those animals were considered to be adults.

**Technical Appendix References**

11. Strauss E, Ronnenberg K, Klages I, Graeber R. Wildlife survey in Lower Saxony 1991-2016 - a base tool for description of biodiversity of our cultural landscape. Verh Ges Oekol. 2016;363-364.

12. Suchentrunk FW, Willig R, Hartl GB. On eye lens weights and other age criteria of the Brown hare *(Lepus europaeus* Pallas, 1778). Z Säugetierkd. 1991;56:365-374.

13. Broekhuizen S, Maaskamp F. Age determination in the European hare (*Lepus europaeus* Pallas) in the Netherlands. Z Säugetierkd. 1979;44(3):162-174.
